# Gain of channel function and modified gating properties in TRPM3 mutants causing intellectual disability and epilepsy

**DOI:** 10.1101/2020.04.02.021758

**Authors:** Evelien Van Hoeymissen, Katharina Held, Ana Cristina Nogueira Freitas, Annelies Janssens, Thomas Voets, Joris Vriens

**Affiliations:** Laboratory of Endometrium, Endometriosis and Reproductive Medicine, Department of Development and Regeneration, KU Leuven, Belgium; Laboratory of Ion Channel Research, VIB-KU Leuven Center for Brain and Disease Research, Leuven, Belgium and Department of Molecular Medicine, KU Leuven, Belgium

**Author notes:** Shared first authors (Evelien Van Hoeymissen, Katharina Held, Ana Cristina Nogueira Freitas). Shared last and corresponding authors (Thomas Voets, Joris Vriens). Corresponding authors: Prof. Joris Vriens, Laboratory of Endometrium, Endometriosis and Reproductive Medicine, Department of Development and Regeneration, KU Leuven, Herestraat 49 box 611, 3000 Leuven, Belgium, Prof. Thomas Voets, Laboratory of ion channel Research, VIB-KU Leuven Center for Brain and Disease Research and KU Leuven Department of Molecular Medicine, Herestraat 49 box 801, 3000 Leuven, Belgium, Phone: +32 16 3.

## Abstract

Developmental and epileptic encephalopathies (DEE) are a heterogeneous group of disorders characterized by epilepsy with comorbid intellectual disability. Recently, two *de novo* heterozygous mutations in the gene encoding TRPM3, a calcium permeable ion channel, were identified as the cause of DEE in eight probands, but the functional consequences of the mutations remained elusive. Here we demonstrate that both mutations (V990M and P1090Q) have distinct effects on TRPM3 gating, including increased basal activity, higher sensitivity to stimulation by the endogenous neurosteroid pregnenolone sulphate (PS) and heat, and altered response to ligand modulation. Most strikingly, the V990M mutation affected the gating of the non-canonical pore of TRPM3, resulting in large inward cation currents via the voltage sensor domain in response to PS stimulation. Taken together, these data indicate that the two DEE mutations in TRPM3 result in a profound gain of channel function, which may lie at the basis of epileptic activity and neurodevelopmental symptoms in the patients.

## Introduction

Transient Receptor Potential (TRP) channel TRPM3 is a calcium-permeable cation channel that can be activated by heat (1) and by a variety of chemical ligands, including the endogenous neurosteroid pregnenolone sulphate (PS) (2). TRPM3 is expressed in a large subset of mouse and human somatosensory neurons, where it is involved in the detection of noxious heat and the development of inflammatory pain (1, 3). Moreover, TRPM3 is expressed in several brain areas, including the choroid plexus, cerebellum, cortex and the hippocampal formation (4-6), but its functional role in these areas is unknown. Recently, two *de novo* substitutions in TRPM3 (V990M and P1090Q) were identified as the cause of intellectual disability and epilepsy in eight probands with developmental and epileptic encephalopathy (DEE) (7). However, the consequences of these mutations on TRPM3 function remained elusive. We here demonstrate that both mutations lead to significant gain-of-channel function, including increased basal activity and higher sensitivity to PS and heat. V990M exhibits further pronounced functional alterations, including anomalous activation of the alternative current through the voltage-sensor domain, reduced sensitivity to receptor-mediated inhibition and calcium-dependent inactivation, and lower sensitivity to block by the anticonvulsant primidone.

## Results & Discussion

When performing Fura-2-based calcium imaging on transiently transfected HEK293T cells, we observed significantly higher intracellular Ca^2+^ concentrations ([Ca^2+^]_i_) in cells expressing the two DEE mutants compared to wild type human TRPM3 (WT) (Figure 1A,B), an effect that was much more pronounced in the V990M mutant. Application of primidone or isosakuranetin, both potent TRPM3 antagonists (8, 9), reduced [Ca^2+^]_i_ in cells expressing WT or mutant TRPM3. In absolute terms, the antagonist-induced reduction in [Ca^2+^]_i_ was the largest for the V990M mutant, yet [Ca^2+^]_i_ did not fully return to the level of cells expressing WT (Figure 1A,C). These results suggest that the DEE mutations lead to increased basal channel activity. In line herewith, whole-cell currents in response to voltage steps revealed increased current amplitudes in cells expressing the DEE mutants compared to WT (Figure 1-figure supplement 1).

**Figure 1:**
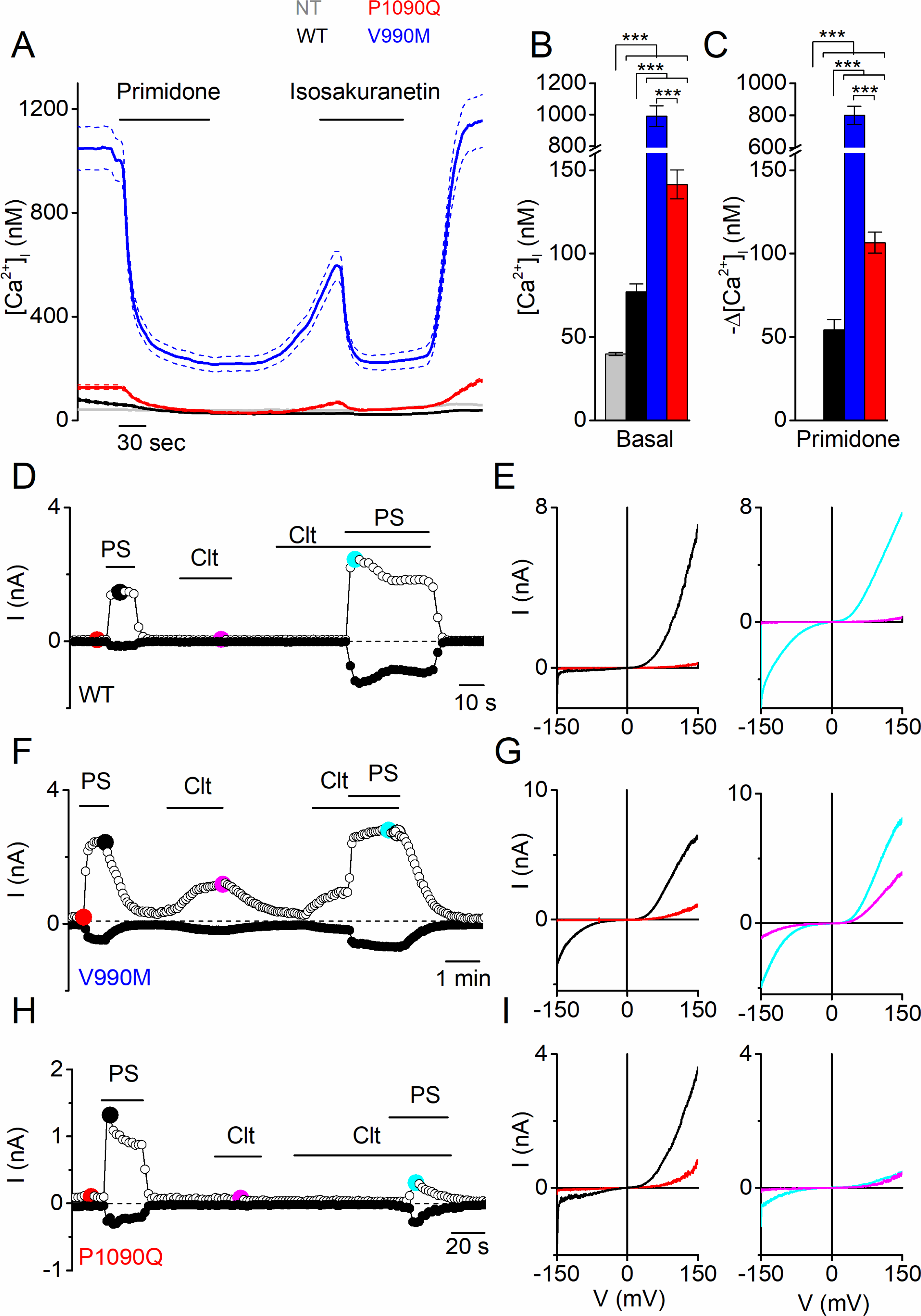
Elevated basal activity in HEK293T cells expressing TRPM3 DEE mutants. (**A**) Time course of intracellular calcium concentrations ([Ca^2+^]_i_) (± SEM) upon application of the TRPM3 inhibitors primidone (100 µM) and isosakuranetin (50 µM) for WT (n = 230), P1090Q (n = 163) and V990M (n = 79) transfected HEK293T cells, and non-transfected (NT) cells (n = 93) (N = 3 independent experiments). (**B**) Basal intracellular calcium concentrations represented as mean ± SEM. (**C**) Primidone-induced decrease in [Ca^2+^]_i_ for the indicated cells. Data are represented as mean ± SEM. (**D, F** and **H**) Amplitude of currents at a holding potential of +80 mV and –80 mV (measured with voltage ramps) upon application of PS (40 µM), Clt (10 µM) and co-application of PS and Clt for WT (n = 10) (**D**), V990M (n = 7) (**F**) and P1090Q (n = 9) (**H**). (**E, G** and **I**) Current-voltage relationships at the time points indicated in (**D**), (**F**) and (**G**). *** p < 0.001 (Kruskal-Wallis ANOVA with Dunn’s posthoc test).

Next, we compared the responses of WT and DEE mutants to stimulation with agonist PS and with clotrimazole (Clt), an antifungal drug and known TRPM3 modulator (10). In line with earlier studies (2), we found that PS (40 µM) reversibly activated outwardly rectifying whole-cell currents in cells expressing WT, whereas application of Clt (10 µM) did not activate currents by itself but potentiated responses to PS (Figure 1D,E and Figure 1-figure supplement 1). In particular, application of PS in the presence of Clt provoked activation of a large inwardly rectifying current component, which has been attributed to activation of an alternative ion permeation pathway located in the voltage-sensing domain, distinct from the central pore (10). The response pattern was strikingly altered in cells expressing the DEE mutants. In the case of V990M, whole-cell currents in response to PS were significantly larger compared to WT, and, notably, exhibited a prominent inwardly rectifying current component. In addition, application of Clt induced robust currents in the absence of PS, whereas currents in the combined presence of Clt and PS were similar in amplitude and shape as WT (Figure 1F,G and Figure 1-figure supplement 1). In the case of P1090Q, whole-cell currents in response to PS were also significantly larger compared to WT, but lacked the inwardly rectifying component observed in the V990M mutant. Clt did not activate currents in cells expressing P1090Q, and, in contrast to WT, inhibited the response to PS (Figure 1H,I and Figure 1-figure supplement 1). Taken together, these results indicate that both DEE mutants lead to significant changes in channel gating, including increased basal activity and pronounced alterations in ligand responses.

To further assess the enhanced response to PS stimulation of the DEE mutants, we compared their apparent affinity to PS by measuring [Ca^2+^]_i_ responses to stepwise increases in PS concentrations. We found that the concentration-response curve for both mutants was shifted to significantly lower concentrations compared to WT. Notably, whereas a PS concentration of 10 µM was required to induce a detectable response in cells expressing WT TRPM3, we observed robust calcium responses at concentrations as low as 100 nM for V990M and 1 µM for P1090Q. Moreover, the maximal increase in [Ca^2+^]_i_ to saturating PS concentrations was significantly higher in P1090Q expressing cells compared to WT (Figure 2A,B). Thus, both DEE mutants show increased responses to the neurosteroid PS.

**Figure 2:**
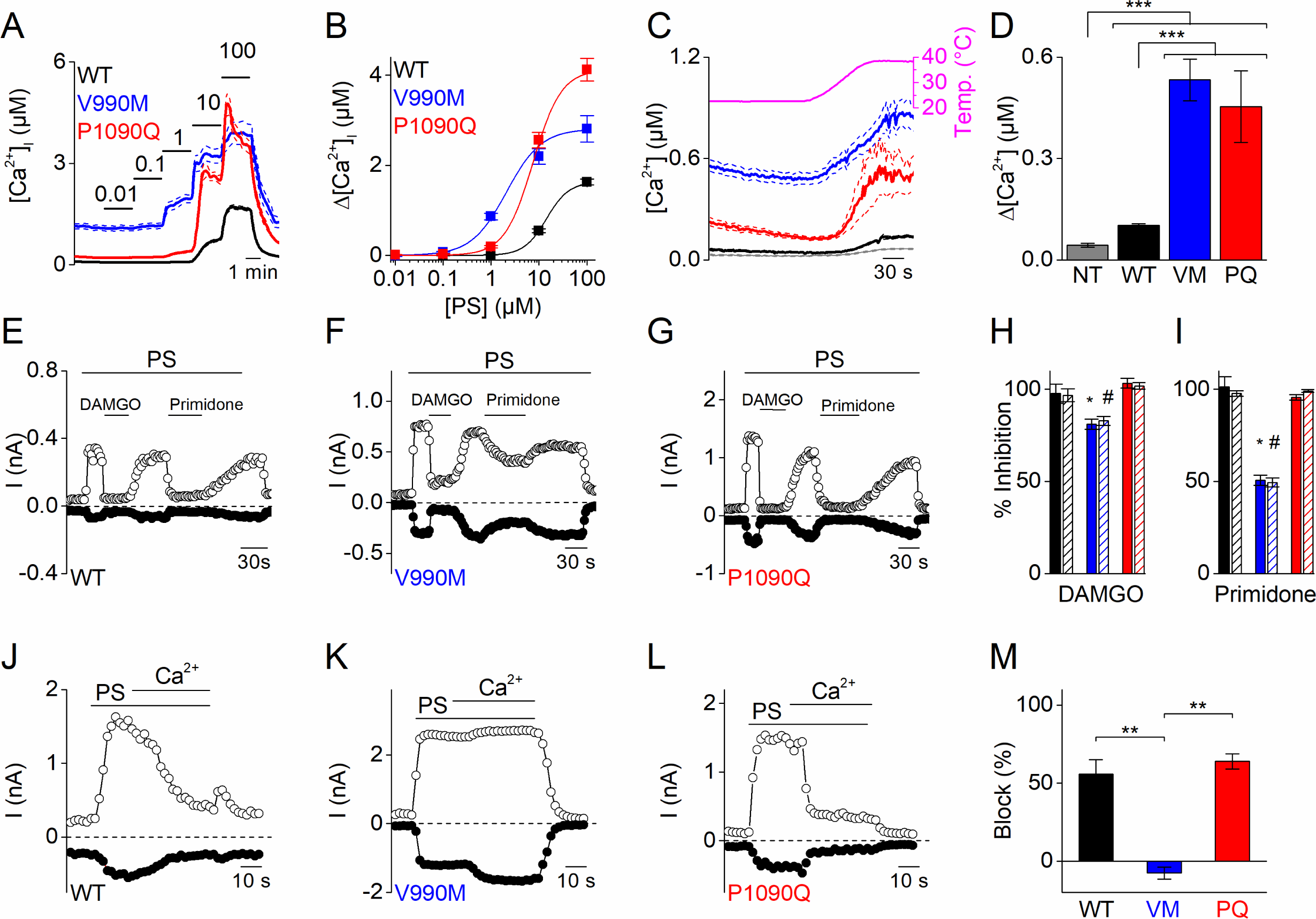
Altered sensitivity of DEE mutants for thermal stimulation and pharmacological modulation. (**A**) Time course of [Ca^2+^]_i_ (± SEM) upon application of the TRPM3 agonist PS in stepwise increasing dose (0.01 – 100 µM) for WT (n = 615), V990M (n = 130) and P1090Q (n = 196) (N = 3 independent experiments). (**B**) PS concentration-response curves for WT (EC_50_ = 14.3 ± 5.8 µM), V990M (EC_50_ = 2.1 ± 0.4 µM), and P1090Q (EC_50_ = 7.5 ± 1.4 µM). (**C**) Time course of [Ca^2+^]_i_ for NT (gray, n= 46), WT (black, n = 148), V990M (blue, n = 259) and P1090Q (red, n = 271) when applying a heat ramp (magenta). Analysis of 3 independent experiments, where the data are represented as mean ± SEM. (**D**) Corresponding amplitudes of the temperature response, represented as mean ± SEM. (**E**-**G**) Amplitude of currents at +80 mV and –80 mV (measured during voltage ramps) upon application of PS (40 µM) with co-application of the µ-opioid receptor agonist DAMGO (1 µM) or the TRPM3 inhibitor primidone (25 µM) for WT (n = 6) (**E**), V990M (n = 8) (**F**) and P1090Q (n = 6) (**G**), in cells co-expressing the µ-opioid receptor. (**H**-**I**) Percentage inhibition of PS-induced currents upon application of DAMGO (**H**) and primidone (**I**) for WT (black), V990M (blue) and P1090Q (red). The filled and shaded bars represent the current inhibition at −80 mV and +80 mV, respectively. (**J**-**L**) Amplitude of currents at +80 mV and –80 mV (measured during voltage ramps) upon application of PS (40 µM) in the presence of 1 mM extracellular calcium for WT (n = 8) (**J**), V990M (n = 6) (**K**) and P1090Q (n = 6) (**L**). (**M**) Percentage inhibition upon calcium application for WT, V990M (VM) and P1090Q (PQ) (mean ± SEM). ** p < 0.01 and *** p < 0.001 (Kruskal-Wallis ANOVA with Dunn’s posthoc test).

TRPM3 is a temperature-sensitive channel, activated upon heating (1). To address whether the DEE mutations affect the channel’s response to heat, we compared changes in [Ca^2+^]_i_ in cells expressing WT or mutant TRPM3 upon exposure to a heat ramp from 23 to 40 °C. Compared to non-transfected cells, we measured a significantly larger increase in [Ca^2+^]_i_ in cells expressing WT or mutant TRPM3. Notably, the amplitude of the heat-induced response was significantly larger in cells expressing the DEE mutants compared to WT (Figure 2C,D).

TRPM3 activity is inhibited upon activation of G protein-coupled receptors, via a mechanism that involves direct binding of the G_βγ_ of trimeric G-proteins to the channel (11-13). To evaluate whether G_β_γ-dependent modulation is altered in the DEE mutations, we co-transfected HEK293T cells with the µ-opioid receptor and WT or mutant TRPM3, and evaluated the effect of the selective agonist DAMGO on PS-activated whole-cell currents. In cells expressing WT or P1090Q, application of 1 µM DAMGO induced a complete and rapidly reversible inhibition of inward and outward currents, whereas V990M was only partly inhibited (Figure 2E-I). The DAMGO concentration for half-maximal inhibition of the PS-activated currents shifted from 4.0 ± 0.6 nM for WT to 40 ± 10 nM for V990M (Figure 2-figure supplement 1). Note that DAMGO was without effect on TRPM3 currents in cells that were not co-transfected with the µ-opioid receptor (Figure 2-figure supplement 1). A difference in sensitivity was also found for the anticonvulsant drug primidone, which at a concentration of 25 µM caused a full inhibition of PS-activated inward and outward currents mediated by WT or P1090Q, but blocked currents mediated by V990M by only ∼50% (Figure 2E-G,I). A similar difference in primidone sensitivity was observed using Fura-2-based calcium imaging (Figure 2-figure supplement 1). When switching from the standard, Ca^2+^-free extracellular solution to a solution containing 1 mM Ca^2+^, PS-activated currents mediated by WT or by P1090Q undergo time-dependent desensitization (10, 14). In contrast, PS-activated currents mediated by V990M remained stable in the presence of extracellular Ca^2+^, indicating reduced sensitivity to Ca^2+^-dependent desensitization in this mutant (Figure 2J-M).

The PS-induced whole-cell currents mediated by V990M showed a prominent inwardly rectifying current component. A similar inwardly rectifying current component can also be activated in WT TRPM3 when PS is applied in the presence of Clt. In earlier work, we have demonstrated that this inwardly rectifying current component represents ion flux through an alternative ion permeation pathway located in the voltage sensor domain of TRPM3, which can be distinguished from the central pore based on its voltage dependence, insensitivity to pore block by La^3+^ and ion selectivity (including a lower permeability for monomethylammonium (MMA^+^) compared to Na^+^) (10, 14). Notably, the V990M mutation is located in close vicinity of Asp988 and Gly991, which we recently identified as critical determinants of the alternative ion permeation pathway (15). We therefore hypothesized that gating of the alternative ion permeation pathway is facilitated in V990M, such that it can be activated by PS even when Clt is not co-applied. We found that the PS-activated current in the V990M mutant showed a bimodal voltage dependence (Figure 3B), its inward current component was resistant to block by the central pore blocker La^3+^ (Figure 3F,G), and inward currents were reduced when extracellular Na^+^ was replaced by MMA^+^ (Figure 3H). Taken together, these data indicate that the V990M mutation leads to a gain of function at the level of the alternative ion permeation pathway.

**Figure 3:**
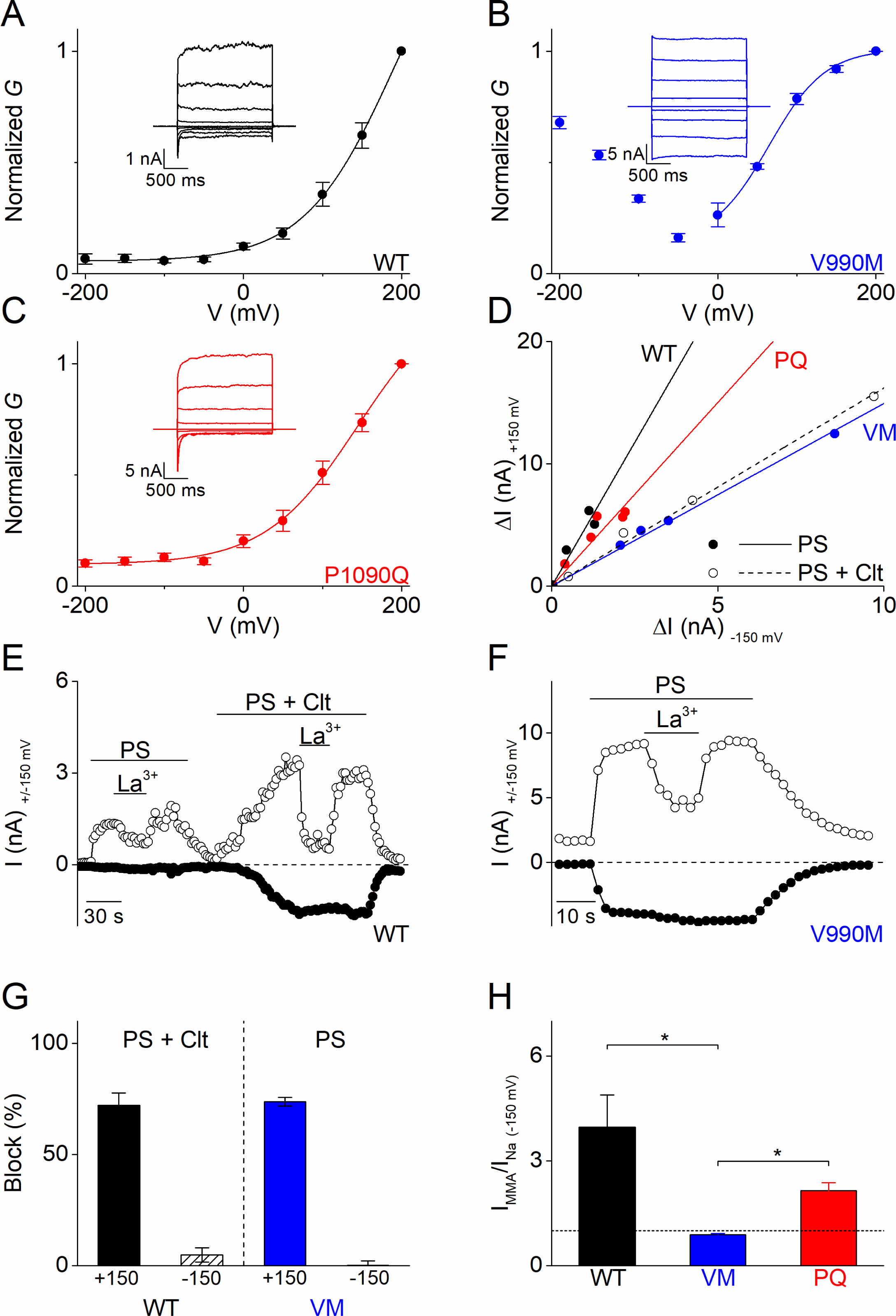
Altered gating of the alternative pore in V990M. **(A**-**C**) *G-V* plots of PS-activated currents for (A) WT (black), (B) V990M (blue) and (C) P1090Q (red). Currents measured during voltage-steps ranging from −200 mV to +200 mV, separated by steps of +50 mV. Representative currents are shown as insets in each graph; n = 6 for each experiment. (**D**) Rectification pattern of PS (40 µM) (full circle and line) and PS + Clt (10 µM)-induced (open circle and dashed line) currents for WT, V990M and P1090Q. Data points are derived by plotting the current increase at +150 mV *versus* the current increases at −150 mV; n ≥ 4 for each dataset. (**E**) Time course of WT TRPM3 whole-cell currents at ±150 mV upon application of PS (40 µM) and Lanthanum (La^3+^; 10 µM) or PS + Clt and La^3+^. (**F**) Time course of V990M mutant whole-cell currents at ±150 mV upon application of PS and La^3+^. (**G**) Relative La^3+^ block calculated from experiments as in E) and F) for WT (black) in presence PS + Clt (n = 4) and for V990M (blue, n = 8) in presence of PS alone. (**H**) Relative PS-induced currents at −150 mV carried by monomethylammonium (MMA^+^) in WT (black), V990M (blue) and P1090Q (red). MMA^+^ currents were normalized to the currents carried by Na^+^; PS (40 µM) and n = 5 for all experiments. ** p < 0.01 (Kruskal-Wallis ANOVA with Dunn’s posthoc test).

Considering that all reported DEE patients were heterozygous for the TRPM3 substitutions (7), and that TRPM3 is functional as a tetramer, it is expected that patients express heteromultimeric channels consisting of variable numbers of WT and mutant subunits. To mimic this situation *in vitro*, we performed a limited number of experiments in cells co-transfected with a mixture of cDNA encoding WT and mutant TRPM3 in a 1:1 ratio. The amplitude of PS-induced inward currents in cells expressing a WT:V990M mixture was intermediate between cells expressing only WT or only V990M (Figure 4A,B). Moreover, the PS-activated currents exhibited the typical inwardly rectifying current component (Figure 4C,D). Finally, in contrast to WT but like V990M, Clt (10 µM) activated robust currents in cells expressing a WT:V990M mixture (Figure 4A,B). Next, the WT:P1090Q co-transfected cells showed PS-induced current amplitudes that were intermediate between WT and P1090Q transfected cells (Figure 4E,F) and showed a shift in the rectification pattern of the PS-induced currents that was different form the homozygote situation (Figure 4G). Moreover, the effect of Clt pre-incubation was different in co-transfected cells. WT:P1090Q cells showed first a potentiation of the PS responses by pre-incubation of Clt, followed by a time-dependent current inhibition. This was in contrast to the block of PS-induced currents by pre-incubation with Clt for the P1090Q mutant in isolation (Figure 4H).

**Figure 4:**
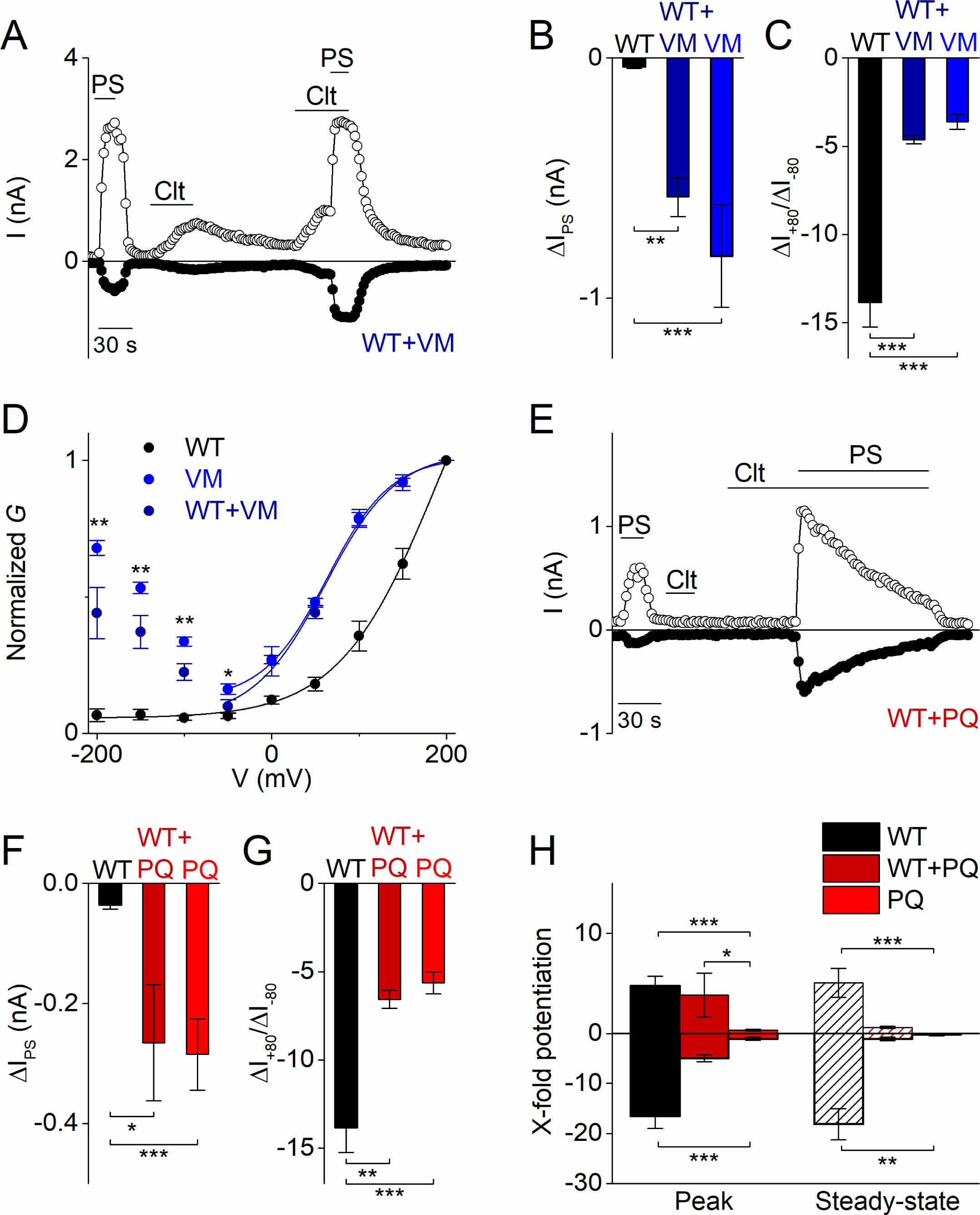
Heterozygous effects of DEE mutants. **(A)** Time course of whole-cell currents at ±80 mV recorded in HEK293T cells transiently co-transfected with WT and V990M mutant DNA (1:1) upon application of PS (40 µM), Clt (10 µM) or PS + Clt. (**B)** Current amplitudes at a holding potential of −80 mV (measured with voltage ramps) upon application of PS (40 µM) for WT (n = 10) (black), VM (n = 7) (light blue) or WT + VM (1:1) (n = 7) (dark blue). (**C)** Same as in (B) but for current amplitude ratios of +80 mV/-80 mV. (**D)** *G-V* plots for PS-activated WT TRPM3 (black), V990M (light blue) and WT+V990M (1:1) (dark blue). Data points were obtained with a step protocol ranging from - 200 mV to +200 mV with +50 mV steps; n ≥ 6 for each experiment. **(E)** Time course of whole-cell currents at ±80 mV recorded in HEK293T cells transiently co-transfected with WT and P1090Q (1:1) upon application of PS (40 µM), Clt (10 µM) or PS + Clt. **(F)** Similar as in (B) but for WT (n = 10) (black), P1090Q (n = 9) (light red) or co-transfected WT + P1090Q (1:1) (n = 5) (dark red). (**G)** Similar as in (C) but for WT (n = 10) (black), P1090Q (n = 9) (light red) or co-transfected WT + P1090Q (1:1) (dark red). (**H)** X-fold potentiation at peak and steady-state conditions of Clt-potentiated PS-currents in WT (black), P1090Q (light red) or co-transfected with WT + P1090Q (1:1) (dark red); n ≥ 4. * p < 0.05, ** p < 0.01 and *** p < 0.001 (Kruskal-Wallis ANOVA with Dunn’s posthoc test for panel B, F and H. One-way ANOVA with Tukey’s posthoc test for panel C and G).

In conclusion, our results indicate that two human mutations in the TRPM3 gene associated with DEE give rise to channels with substantially altered functional properties. Whereas the V990M and P1090Q mutations have various differential effects on several aspects of TRPM3 gating, both can be considered as strong gain-of-function mutants, with increased inward cation currents and Ca^2+^ influx under basal condition or when stimulated with heat or the endogenous neurosteroid PS. We hypothesize that the increased calcium influx and depolarizing channel activity may lie at the basis of seizure development and neurodevelopmental symptoms in DEE patients.

## Acknowledgements

We thank all the members of the Laboratory of Ion Channel Research and the Laboratory of Endometrium, Endometriosis and Reproductive Medicine at the KU Leuven, for their helpful discussions and comments.

## Figure legends

**Figure 1-figure supplement 1:**
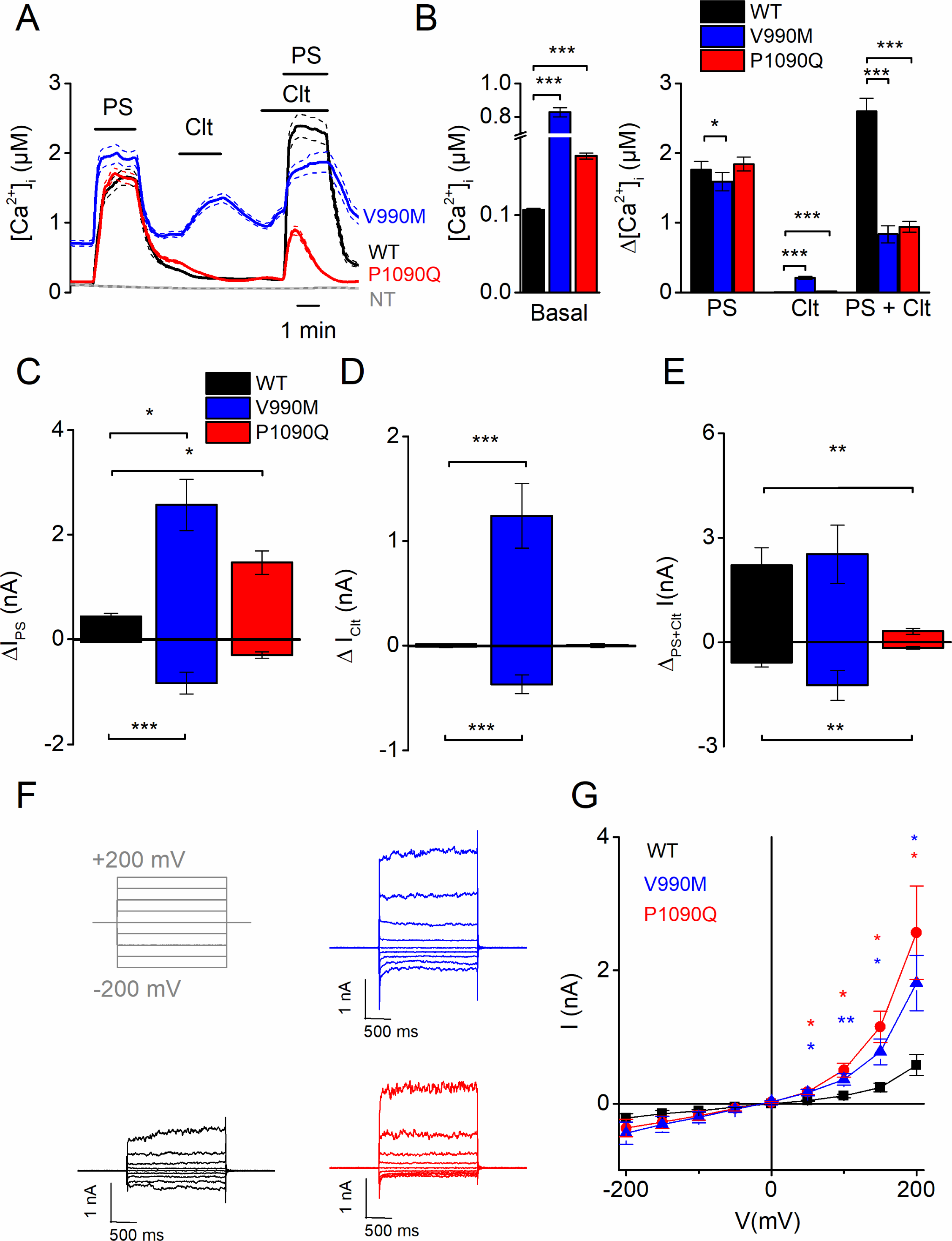
Biophysical characterization of the V990M and P1090Q substitution in hTRPM3 indicate the substitutions as a gain of function mutation. **(A)** Time course of calcium concentrations (± SEM) upon application of PS (40 µM), Clt (10 µM) and co-application of PS and Clt for WT (n = 294), V990M (n = 196) and P1090Q (n = 624) transfected cells. (**B**) Basal [Ca^2+^]_i_ (left) and calcium amplitudes when applying PS, Clt and co-application of PS and Clt (right) for WT, V990M and P1090Q transfected cells. (**C**-**E**) Amplitude of currents at a holding potential of + 80 mV and – 80 mV (measured with voltage ramps) upon application of PS (40 µM) (**C**), Clt (10 µM) (**D**) and co-application of PS and Clt (**E**). (**F**) Representative whole-cell TRPM3 currents recorded during voltage steps ranging from −200 mV to +200 mV, separated by steps of +50mV for WT (black), V990M (blue) and P1090Q (red) transfected cells. (**G**) Current-voltage relationship for WT (n = 5), V990M (n = 6) and P1090Q (n = 7) transfected cells. * p < 0.05, ** p < 0.01 and *** p < 0.001 (Kruskal-Wallis ANOVA with Dunn’s posthoc test for panel B-E. Two-way ANOVA with Tukey’s posthoc test for panel G). Data are represented as mean ± SEM.

**Figure 2-figure supplement 1:**
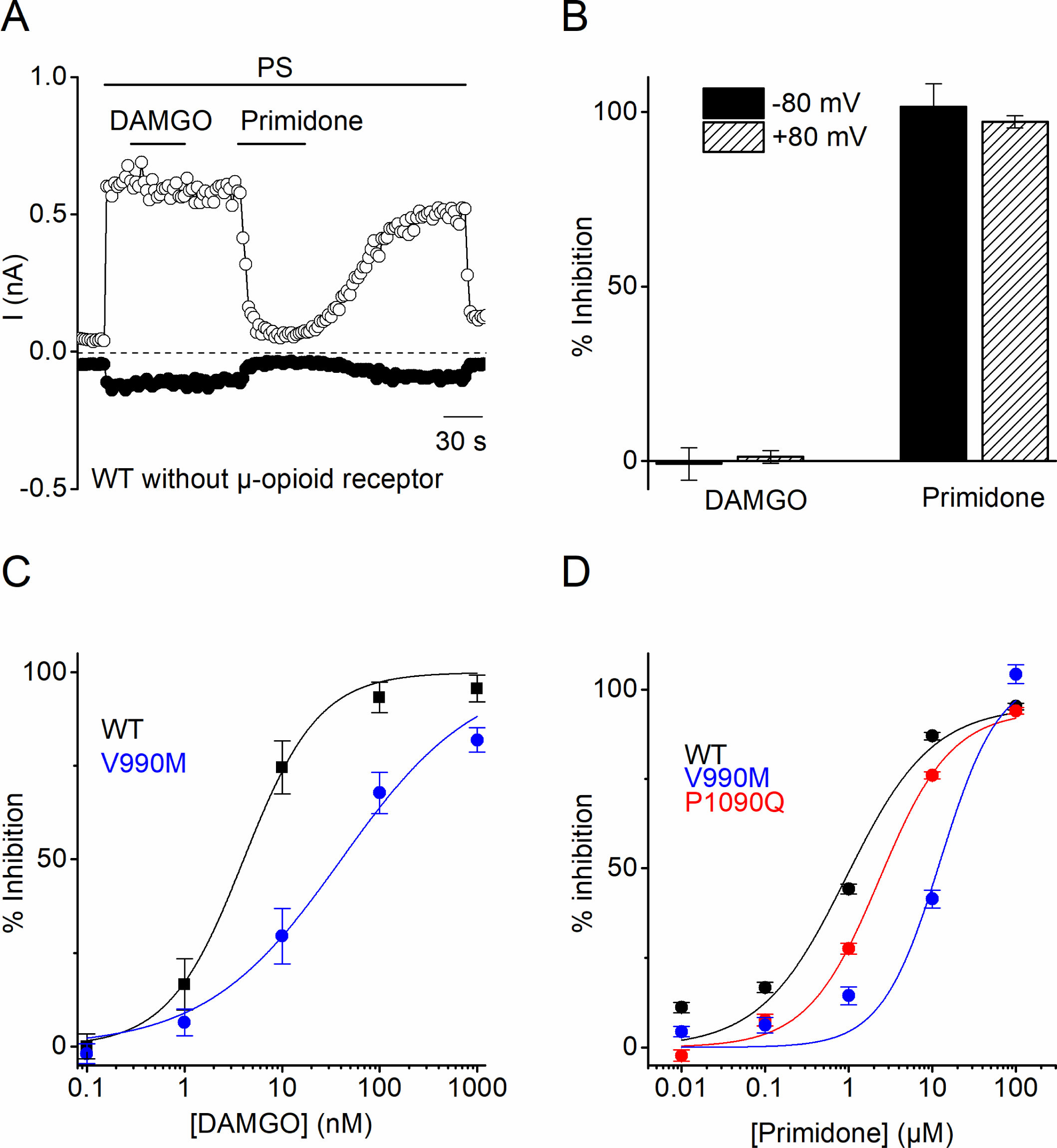
Characterization of DAMGO and primidone block on PS-induced currents. (**A**) Amplitude of currents at a holding potential of + 80 mV and – 80 mV (measured with voltage ramps) upon application of PS (40 µM) with co-application of the µ-opioid receptor agonist DAMGO (1 µM) and the TRPM3 inhibitor primidone (25 µM) for WT hTRPM3 transfected cells (n = 5). (**B**) Percentage inhibition of PS-induced currents at holding potential of + 80 mV and – 80 mV upon application of DAMGO and primidone for WT hTRPM3 cells. (**C**) DAMGO concentration-response curve for WT (IC_50_ = 4.0 ± 0.6 nM) and V990M (IC_50_ = 40 ± 10 nM) hTRPM3 co-transfected cells with µ-opioid receptors (n = 5). (**D**) Primidone concentration-response curve for WT (IC_50_ = 940 ± 250 nM; n = 526), V990M (IC_50_ = 12.5 ± 3.7 µM; n = 498) and P1090Q (IC_50_ = 2.3 ± 0.3 µM; n = 538) hTRPM3 transfected cells (N = 3 independent calcium imaging experiments).

## Supplementary Section

### Material & Methods

#### Cell culture

HEK293T cells were cultured as described previously (15). HEK293T cells were transiently transfected with 2 µg of DNA using TransIT transfection reagent (Mirus) 36-48 hours before the measurements. In all co-transfection experiments a ratio of 1:1 between WT and DEE mutant DNA was used, to end up with a total of 2 µg of DNA.

#### Site-directed Mutagenesis

All mutants were obtained by the standard PCR overlap extension method using hTRPM3 directly linked to YFP from pCAGGS/IRES-GFP vector (15). Accuracy of all mutant sequences was verified by sequencing of the entire DNA constructs.

#### Fluorescence imaging

Changes in intracellular calcium concentration were monitored using ratiometric Fura-2-based fluorimetry. Cells were loaded with 2 µM Fura-2-acetoxymethyl ester (Alexis Biochemicals) for 30 min at 37 °C. Fluorescence was measured during alternating illumination at 340 and 380 nm using Eclipse Ti (Nikon) fluorescence microscopy system, and absolute calcium concentration was calculated from the ratio of the fluorescence signals at these two wavelengths (R = F_340_/F_380_) as [Ca^2+^] = K_m_ × (R-R_min_)/(R_max_-R), where K_m_, R_min_ and R_max_ were estimated from in vitro calibration experiments with known calcium concentrations. The bath solution contained (in mM) 138 NaCl, 5.4 KCl, 2 CaCl_2_, 2 MgCl_2_, 10 glucose, and 10 HEPES, pH 7.4. Pregnenolone sulphate, and clotrimazole were obtained from Sigma-Aldrich and dissolved in bath solution from a 1000x stock solution in DMSO.

#### Whole cell patch clamp recordings

Standard whole-cell patch-clamp recordings were performed with an EPC-10 amplifier and the PatchMasterPro Software (HEKA Elektronik, Lambrecht, Germany). Current measurements were performed at a sampling rate of 20 kHz and currents were digitally filtered at 2.9 kHz. In all measurements, 70% of the series resistance was compensated. The standard internal solution contained (in mM): 100 CsAsp, 45 CsCl, 10 EGTA, 10 HEPES, 1 MgCl_2_ (pH 7.2 with CsOH) or 140 potassium gluconate, 5 EGTA, 1 MgCl_2_, 10 HEPES and 2 NaATP (pH 7.3 with CsOH) for the measurements of G_βγ_ mediated inhibition and the standard extracellular solution contained (in mM): 150 NaCl, 1 MgCl_2_, 10 HEPES (pH 7.4 with NaOH). The standard patch pipette resistance was between 2 MΩ and 4 MΩ when filled with pipette solution. In experiments allowing Ca^2+^-dependent desensitization of TRPM3, Mg^2+^ was replaced by Ca^2+^ in the extracellular solution, and Cs^+^ was replaced by Na^+^ in the pipette solution.

#### Statistics

Electrophysiological data were analyzed using IgorPro 6.2 (WaveMetrics, USA), WinASCD (Guy Droogmans, Leuven) and OriginPro 8.6 (OriginLab Corporation, USA). OriginPro 8.6 was further used for statistical analysis and data display. For the statistical comparison of data sets, we first tested for normality using the Shapiro-Wilk test and for homogeneity of variance using Levene’s test. Based on the outcome of these tests, we used either ANOVA followed by Tukey’s posthoc test (when there was no significant variance inhomogeneity and the data were significantly drawn from a normal distribution) or Kruskal-Wallis ANOVA with Dunn’s posthoc test (when the requirement of similar variance or normal distribution were not met). Posthoc tests were only performed when the overall ANOVA or Kruskal-Wallis ANOVA revealed overall significance. Used tests are noted in the figure legends. P values below 0.05 were considered as significant. Data points represent means ± SEM of the given number (n) of identical experiments.

Conductance-voltage (*G-V*) curves were fitted with a Boltzmann function of the form:

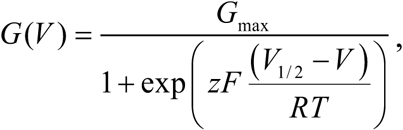

where *z* is the apparent gating charge, *V*_*1/2*_ the potential for half-maximal activation, *G*_*max*_ the maximal conductance, *F* the Faraday constant, *R* the gas constant and *T* the absolute temperature. Experiments were performed at room temperature.

